# Auditory brainstem-cortical anatomy constrains the magnitude of frequency-following responses (FFRs) and event-related potentials (ERPs) coding speech-in-noise

**DOI:** 10.64898/2026.01.03.697410

**Authors:** Gavin M. Bidelman, Jack R. Stirn, Rose Rizzi, Jessica A. MacLean, Hu Cheng

## Abstract

Speech-evoked brain potentials provide a window into the neural encoding of speech, experience-dependent plasticity, and deficits in central auditory processing from communication disorders. Stronger and faster frequency-following responses (FFRs) and cortical event-related potentials (ERPs) have been interpreted as reflecting more robust and efficient auditory-sensory processing across brainstem and cortical levels. Importantly, these neural signatures relate to real-world listening skills like speech-in-noise (SIN) perception. Yet, how these speech-evoked FFRs/ERPs relate to underlying auditory anatomical structures is unknown. Using a multimodal imaging approach, we recorded FFRs and ERPs to clean and noise-degraded speech sounds to assess the strength of listeners’ neural encoding of speech at brainstem (FFR) and cortical (ERP) levels. MRI volumetrics of midbrain and transverse temporal gyrus (Heschl’s gyrus) quantified morphological variation in subcortical and cortical anatomy that underly these EEG potentials. The QuickSIN assessed behavioral SIN abilities. Results showed that larger and thicker right (but not left) Heschl’s gyrus was related to listeners’ SIN abilities as well as the size of their cortical ERPs. Structural and functional measures interacted at a subcortical level. For listeners with smaller midbrain volumes, larger speech FFRs were associated with better QuickSIN scores; whereas in individuals with larger midbrain volumes, larger FFRs were related to poorer QuickSIN. Our findings reveal common functional signatures of speech processing (FFRs, ERPs) are constrained by the anatomy of their underlying generators and suggest a complex interplay between auditory brain structure and function in accounting for perceptual SIN capacity.

## 1. Introduction

Everyday listening situations contain noise that hinders speech perception [1]. Issues with speech-in-noise (SIN) understanding are familiar complaints among the elderly and listeners with hearing loss [2] but are also common in individuals with unremarkable audiological findings [3-7]. Perceptual figure-ground deficits are also observed in many language and auditory-based disorders despite normal audiograms [8-13]. As such, it is now well recognized that SIN listening skills are determined by more than audibility or peripheral hearing status [1,3-5,14,15]. SIN tests are now among the most informative assessments of real-world listening capacity [16,17]. This has led several investigators to examine whether electrophysiological assays might provide a window into how the central auditory nervous system encodes speech sounds, and in turn, whether certain EEG biomarkers might predict SIN skills at the individual level [3,5,15,18].

The brain’s neuroelectric response to speech reflects an aggregate of EEG activity generated from both brainstem and cortical structures [19]. The cortical event-related potentials (ERPs) are composed of several waves (e.g., P1-N1-P2), reflecting activation of auditory thalamus, cortex, and associative areas [20]. The brainstem component, termed the frequency-following response (FFR), is a sustained potential that mirrors acoustic stimuli with high fidelity [21-23]. Neural generators of the FFR have been described in previous M/EEG studies [4,23-27]. And while they reflect a mixture of phase-locked activity from structures throughout the auditory pathways [23-25,28], FFRs are dominantly of brainstem (midbrain) origin when recorded to sounds with frequencies >150 Hz [4,23,24]. Links between noise-related changes in both brainstem (FFR) and cortical (ERP) speech-evoked potentials and SIN perception have been widely reported over the past decade [e.g., 3,4-6,15,18,29-35]. In general, faster and more robust FFRs/ERPs are associated with better neural representation of speech and consequently, better speech perception in noise-degraded listening tasks. Collectively, a growing body of studies suggest speech-FFRs/ERPs are valuable neurophysiological correlates for tracking both positive and maladaptive auditory plasticity in the brain related to speech and SIN perception.

Despite their potential as an objective “barometer” of complex auditory processing [36], there is a surprising dearth of studies examining links between these functional EEG markers and the underlying brain anatomy from which they originate. Presumably, morphology of auditory structures should constrain their corresponding evoked potentials. Given the basic principles of electrophysiology and volume conduction [37,38], larger size (volume, thickness, area) of the brainstem-cortical auditory pathways should support increased neural synchrony and packing of the neural elements generating electrical brain potentials like FFRs and ERPs.

In this vein, a handful of multimodal imaging studies combining M/EEG and MRI have shown connections between the morphology and neurophysiology of Heschl’s gyrus (HG) and surrounding auditory cortex. For example, reduced volume of HG is observed in patients with tinnitus [39] while larger HG volumes are observed in musicians [40] and listeners with absolute pitch [41]. In particular, Schneider, *et al*. [40] reported evoked MEG responses from primary auditory cortex were 102% larger and gray matter volume 130% larger in professional musicians. In older adults, subtle declines in gray matter volume of primary auditory cortex accounts for reduced hearing sensitivity [42]. These studies establish structure-function relationships at a *cortical* level of the auditory system. However, only a handful of studies have assessed, albeit indirectly, auditory structure-function relations at the *brainstem* level.

One study showed that hemodynamic activation in right auditory cortex, as measured via fMRI BOLD signal, is related to individual differences in the strength of cortically-based FFRs [43]. Microstructural integrity of the white matter surrounding left primary auditory areas also related to FFR latency, suggesting a link between the fine timing of neural phase-locked responses and left lateralized auditory brain networks [43]. Relatedly, fMRI activation in the midbrain (i.e., inferior colliculus) predicts FFR changes in pitch tracking accuracy of voice fundamental frequency (F0) after auditory learning [44]. While these studies reveal correspondences between different neuroimaging assays, we are unaware of any study that has assessed whether volumetric properties of auditory midbrain and cortical anatomy map to functional properties of speech-evoked FFRs and ERPs, respectively.

To this end, we used a multimodal imaging approach to record FFRs and ERPs to clean and noise-degraded speech sounds via EEG to evaluate the strength of listeners’ neural encoding of SIN. MRI volumetrics of the midbrain and HG derived from anatomical MRIs provided insight into morphological variation in subcortical and cortical auditory brain anatomy that underly speech-FFR/ERP responses. We hypothesized physical differences in auditory brainstem and cortex would have measurable associations with FFR and ERP magnitudes. Our findings reveal that the strength of listeners’ electrophysiological responses to speech is partially constrained by their underlying auditory brain anatomy.

## 2. Materials and Methods

### 2.1 Participants

We collected EEG and MRI data from N=30 adults (6 male, 24 female) aged 18-41 years (μ ± σ = 23.3 ± 4.7 years). Participants were recruited from the Indiana University student body and Greater Bloomington area. All exhibited normal hearing sensitivity confirmed via audiometric screening (i.e., < 25 dB HL, octave frequencies 250 -8000 Hz). Most were right-handed (mean laterality index; 74.3±40.7) [45]. All had a collegiate level of education (16.6 ± 3.0 years formal education) and were native speakers of American English. On average, the sample had obtained 8.6 ± 7.9 years (range = 0 – 35 years) of formal, self-reported music training. All were paid for their time and gave informed consent in compliance with a protocol approved by the Institutional Review Board at Indiana University.

### 2.2 EEG recording and analysis

We recorded brainstem (FFRs) and cortical (ERPs) evoked potentials simultaneously as participants monitored a triplet of vowel sounds (/a/, /i/,/u/) presented continuously for ∼15 min. totaling ∼2000 trials of each token [for full EEG task details, see 18]. Occasional deviants (/u/ vowels; 70 trials) were pseudo-randomly intermixed within the vowel stream. Participants were asked to detect the oddball sounds via a key press. Each vowel was 100 ms with a common voice F0 (=150 Hz). This F0 is above the phase-locking limit of cortical neurons, ensuring our FFRs would be primarily of brainstem origin [46,47]. Auditory stimuli were delivered at 75 dBA SPL through shielded [18] ER-2 insert headphones (Etymotic Research, Elk Grove Village, IL). In addition to the clean run, the task was performed in noise where vowel stimuli were mixed with 8 talker babble (cf. Killion et al., 2004) at a signal-to-noise ratio (SNR) of +5 dB (speech at 75 dBA SPL; noise at 70 dBA SPL).

EEG was recorded differentially between Ag/AgCl disc electrodes placed at the scalp vertex referenced to linked mastoids (M1/M2; mid-forehead = ground). This single-channel montage is ideal for simultaneously recording brainstem and cortical auditory responses which are distributed maximally over frontocentral scalp locations [26,32,48-51]. Interelectrode electrode impedances were ≤ 5 kΩ. EEGs were digitized at 5 kHz (Compumedics Neuroscan, SynAmps RT amplifiers) and corrected for ocular artifacts^1^. Cleaned EEGs were epoched (-10–200 ms), pre-stimulus baselined, and ensemble averaged across trials to obtain compound speech-evoked potentials [19]. We bandpass filtered full-band responses from 100 to 1500 Hz and 1 to 30 Hz to isolate the FFRs and ERPs, respectively [18,19,51].

From FFRs, we measured the RMS amplitude from the steady-state portion (10–100 ms) of the response waveform to quantify the overall strength of the brainstem response to speech (see **Fig. 2b**). From the ERPs, we measured the peak-to-peak amplitude of the N1-P2 complex (see **Fig. 2a**). N1 was identified as the greatest negative deflection between 90 and 145 ms; and P2 as the maximum positive deflection within 145–175 ms [18]. Together, FFR_rms_ and N1-P2 magnitude describe the overall strength of the FFR and ERP and allowed us to directly compare the magnitude of speech encoding at brainstem vs. cortical levels [18,54,55].

### 2.3 Structural MRI and morphometric analysis

We acquired a 3D whole-brain T1-weighted anatomical image for each participant using the MPRAGE sequence (TR/TE = 2400/2.68 ms; field of view = 256 × 240 mm^2^; 208 sagittal slices; voxel size = 0.8 mm isotropic; flip angle = 8°) using the Siemens 3T Magnetom Prisma Fit scanner at the IU Imaging Research Facility. MRI data were converted to Brain Imaging Data Structure (BIDS) format using ezBIDS (https://brainlife.io/ezbids/) for further processing [56].

Cortical reconstruction and volumetric segmentation were performed on the T1-weighted structural MRI scans using the FreeSurfer (v7.3.1; http://surfer.nmr.mgh.harvard.edu/) comprehensive recon-all pipeline [57]. This automated pipeline includes motion correction [58], removal of non-brain tissue and scull stripping via hybrid watershed/surface deformation [59], automated Talairach transformation, segmentation of white and gray matter volumetric structures [59,60], and cortical surface reconstruction [61]. Parcellation of brainstem structures (medulla oblongata, pons, midbrain and superior cerebellar peduncle) was performed using FreeSurfer’s Bayesian brainstem segmentation pipeline [62]. MRI processing was conducted on the Indiana University high-throughput Quartz supercomputing cluster (92 compute nodes, each equipped with two 64-core AMD EPYC 7742 2.25 GHz CPUs and 512 GB of RAM).

From the full-brain FreeSurfer output (i.e., aparc + aseg stats table from recon-all), we measured cortical thickness (mm), surface area (mm^2^), and gray matter volume (mm^3^) [59,61,63,64] from each region defined in the Desikan-Killiany atlas parcellation [60]. FreeSurfer morphometrics have good test-retest reliability across scanners and various field strengths [65,66]. Statistical results were projected onto the cortical surface using FreeSurfer’s *fsaverage* standard brain [67].

### 2.4 QuickSIN speech-in-noise perception task

We measured listeners’ SIN perception using the QuickSIN [68]. Participants heard six sentences embedded in four-talker noise babble, each containing five keywords. Sentences were presented at 70 dB HL. The SNR decreased parametrically in 5 dB steps from 25 dB SNR to 0 dB SNR. At each SNR, participants were instructed to repeat the sentence, and correctly recalled keywords were logged. We computed their SNR loss by subtracting the number of recalled target words from 25.5 (i.e., SNR loss = 25.5-Total Correct). The QuickSIN was presented binaurally via Sennheiser HD 280 circumaural headphones. Two lists were run and the second was used in subsequent analysis to avoid familiarization effects [69,70].

### 2.5 Statistical analysis

Unless otherwise noted, we analyzed the dependent variables using linear models in R (v4.2.2) [71]. Subjects served as a random effect for repeated measures models. QuickSIN scores served as the main dependent variable. Subject demographics (e.g., sex, age, music training) were unrelated to QuickSIN scores (all *p*s > 0.44) so these were not included in the models. Tukey-adjusted contrasts were used for multiple comparisons. Visualization of morphometric statistics was performed in the *fsbrain* package [72] and Freeview [57].

## 3. Results

### 3.1 Behavioral results

**Figure 1a** shows QuickSIN scores across the sample plotted against listeners years of formal music training and age. Neither age, music, sex, nor pure-tone average (PTA) hearing levels were associated with associated with QuickSIN scores (all *p*s > 0.07). However, despite normal hearing acuity in our sample (**Fig. 1b**), QuickSIN scores showed substantial variability across listeners (range = -2.5 to 3.5 dB SNR loss) confirming large variation in SIN performance even in young, normal-hearing adults [e.g., 3,4,5,18].

**Figure 1:**
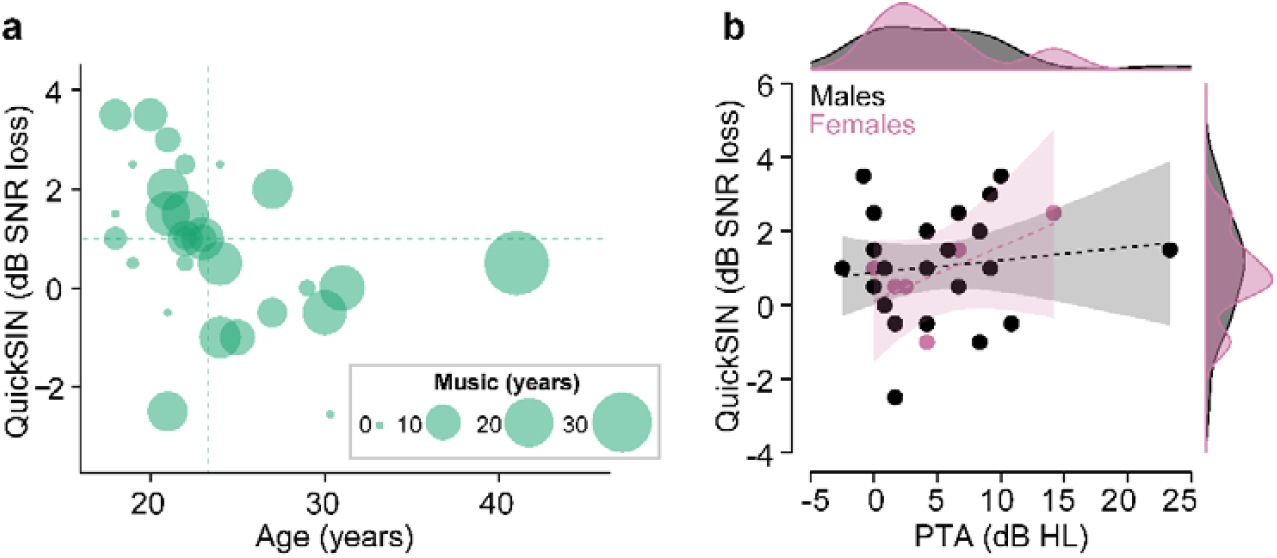
Behavioral speech-in-noise performance. (**a**) QuickSIN scores reflect signal-to-noise ratio thresholds for sentence recognition in noise [68]. QuickSIN scores are plotted against listeners’ age and music training (which were not related to SIN performance). Dotted lines=variable means. (**b**) Plots of hearing acuity (PTA) and QuickSIN scores as a function of sex. Males and females had similar SIN performance and PTA. Clinically “normal” hearing is ≤ 25 dB HL. Dotted lines= *n*.*s*. regression. shading = 95% CI.

### 3.2 Speech-evoked EEG results

Figure 2. shows cortical ERPs and brainstem FFRs as a function of noise. Cortical ERPs showed a series of biphasic deflections (“P1-N1-P2” waves) over the first ∼200 ms after sound onset, consistent with typical morphology of the canonical auditory evoked potentials (**Fig. 2a**). The N1-P2 complex was maximal near the scalp vertex and inverted at the mastoids, consistent with generators in the supratemporal plane (e.g., auditory cortex) [20,73,74]. FFRs appeared as a phase-locked neurophonic potential that mirrored spectrotemporal properties of the eliciting speech stimulus (**Fig. 2b**). FFR spectra revealed that subcortical activity captured the voice fundamental frequency (F0) and its integer related partials up to about the 8^th^ harmonic (∼1200 Hz), consistent with the upper limit of phase locking in human midbrain [75].

Noise weakened and prolonged both the FFRs and ERPs suggesting poorer neural representation of speech at both sub- and neo-cortical levels of auditory processing. Comparison between SNRs revealed noise-related decline in both brainstem FFR [clean vs. noise: *t*(29) = 2.14, *p* = 0.041] and cortical ERP [clean vs. noise: *t*(29) = 4.32, *p* = 0.00017] amplitudes. A linear model revealed behavioral QuickSIN scores were critically dependent on both FFR and ERP strength [FFR_RMS_ x N1-P2 interaction: *F*(1,26) = 4.50, *p* = 0.0049]. As these EEG effects replicate many prior studies examining noise-related changes in the speech-FFRs/ERPs and their relation to SIN perception in both young and older adults [5,6,18,32,76-79], we did not pursue electrophysiological effects further. Instead, we focused subsequent analyses on how structural properties of the brainstem-cortical auditory anatomy link to functional (FFR/ERP) and behavior assays of SIN processing.

**Figure 2:**
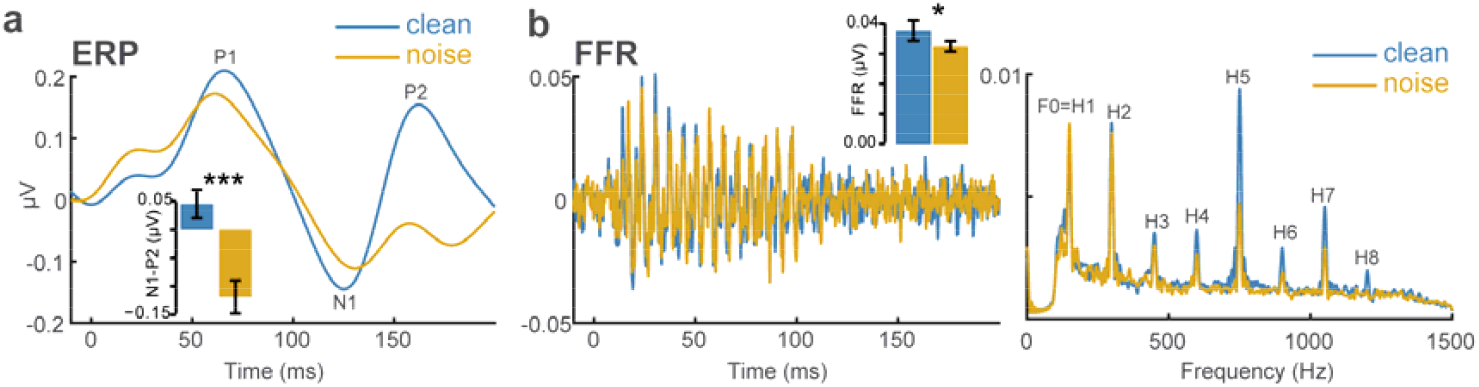
Grand average ERPs and FFRs show noise-related changes in the neural encoding of speech. (**a**) Cortical ERPs. (**b**) Brainstem FFR time waveforms (*left*) and response spectrum (*right*). Waveforms reflect activity at the Cz electrode (only vowel /a/ responses shown for clarity). Noise decreases amplitude and prolongs latency of neural responses including the N1-P2 of the ERPs and RMS amplitude of the FFR (inset bar charts). F0 = fundamental frequency; H1-H8 = harmonics. errorbars = ±1 s.e.m. \**p* < 0.05, \*\*\**p* < 0.0001.

### 3.3 Structural MRI results

Structural morphology of auditory cortex (HG) was related to SIN perception but depended on hemisphere (**Fig. 3**).

**Figure 3:**
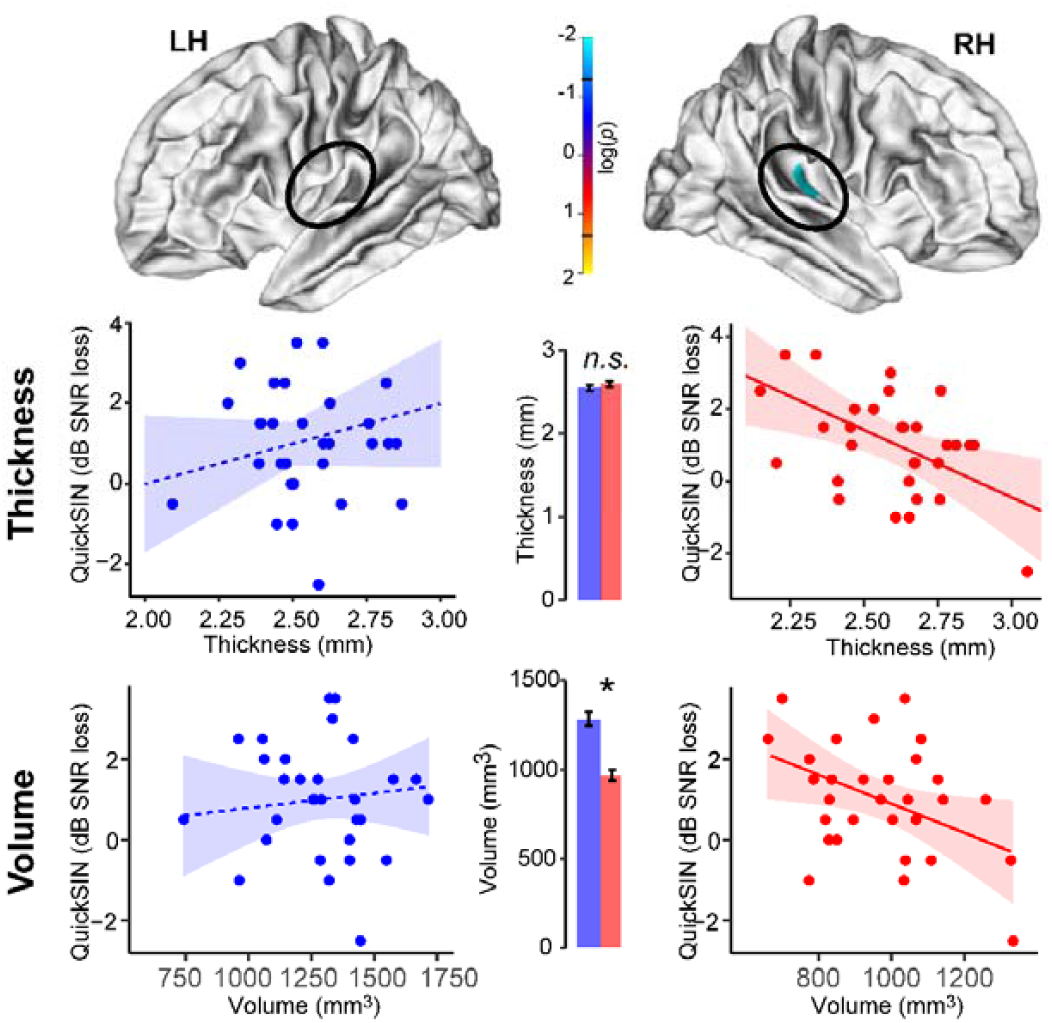
Structural asymmetries in auditory cortex relate to SIN perception. (*top*) Statistical maps show the correlation between tissue thickness and QuickSIN scores projected onto the cortical surface (i.e., FreeSurfer, *mri_glmfit)* [57]. Brain images are masked for the HG region of interest (oval) according to the DK atlas [60]. (*middle*) Cortical thickness in right (but not left) HG predicted QuickSIN performance. (*bottom*) HG volume was smaller in right vs. left hemisphere (inset). As with thickness, more voluminous right HG was associated with better QuickSIN scores. Dotted lines = *n*.*s*. correlations; solid lines = significant correlations. shading = 95% CI. *\*p* < 0.05.

#### 3.3.1 Cortical thickness

Cortical thickness of transverse HG did not differ between right and left hemisphere [*t*(29) = -1.22, *p* = 0.23]. However, a linear model revealed that right [*F*(1,27) = 9.38, *p* = 0.0049] but not left [*F*(1,27) = 0.019, *p* = 0.89] auditory cortex thickness was associated with QuickSIN scores (Fig. 3, *middle panels*). Thicker right HG was associated with lower QuickSIN scores indicative of better SIN performance.

#### 3.3.2 Cortical volume

In contrast to thickness, cortical volume was larger in the left vs. right HG [*t*(29) = 8.25, *p*< 0.0001], consistent with well-known leftward asymmetry and larger density of auditory cortex [80-82]. Paralleling cortical thickness, we found that right [*F*(1,27) = 9.38, *p* = 0.0049] but not left [*F*(1,27) =0.019, *p* = 0.89] auditory cortex volume was also associated with QuickSIN scores (Fig. 3, *bottom panels*). More voluminous right HG was associated with better SIN performance.

#### 3.3.3 Control analysis

We also examined whether QuickSIN scores varied with morphology of adjacent primary motor regions [42], given the putative role of motor system in SIN processing [83]. These control analyses showed that morphology of precentral gyrus in either hemisphere was unrelated to QuickSIN scores (all *p*s > 0.19). This confirms that SIN perception was driven by morphology specific to auditory cortex rather than gross differences in brain anatomy.

#### 3.3.4 Brainstem volumetrics

Volumetric data for the brainstem midbrain are shown in **Figure** Left and right auditory cortical volumes are shown for comparison^2^. A linear mixed-model conducted on volumes [i.e., vol ∼ roi+(1|sub)] revealed a main effect of region [F(2,87) = 2233.1, p < 0.0001]. Tukey contrasts revealed midbrain volumes were (expectedly) larger than either auditory cortex (both ps < 0.0001). Left HG (1287 mm^3^) was 1.3x larger than right HG (971 mm^3^) (p = 0.0021).

### 3.4 Structure-function-behavior relations underlying SIN processing

Figure 5. shows associations between auditory brain structure (MRI volumetrics; see Fig. 4), function (FFR/ERPs; see Fig. 2), and behavioral SIN perception (QuickSIN; Fig. 1). At a cortical level, a multivariate linear model revealed that right HG volume^3^ [*F*(1,26) =6.75, *p* = 0.015] and N1-P2 amplitude of the speech ERPs [*F*(1,26) = 10.57, *p* = 0.0032] independently predicted QuickSIN scores with no interaction [*F*(1,26) = 0.004, *p*=0.95] (**Fig. 5a**).

In contrast to cortex, brainstem data revealed an interaction between midbrain anatomical size, FFR magnitude, and behavior [*F*(1,26) = 4.14, *p* = 0.05]. To parse this interaction of three continuous variables, we performed a median split of the sample, dividing listeners based on whether they had smaller (lower 50^th^ percentile) or larger (upper 50^th^ percentile) midbrain volumes. The direction of brain-behavior correspondence differed within each of these groups. For listeners with smaller brainstem anatomy, larger FFR_RMS_ amplitudes to noise-degraded speech were associated with better (i.e., lower) QuickSIN scores (β_*std*_ *=* -0.11; **Fig. 5b**). In contrast, in listeners with larger anatomy, larger FFR_RMS_ was associated with poorer (i.e., larger) QuickSIN scores (β_*std*_ *=* 0.37). Thus, behavioral SIN processing depended on a complex interaction between anatomical and functional capacity of auditory midbrain structures.

**Figure 4:**
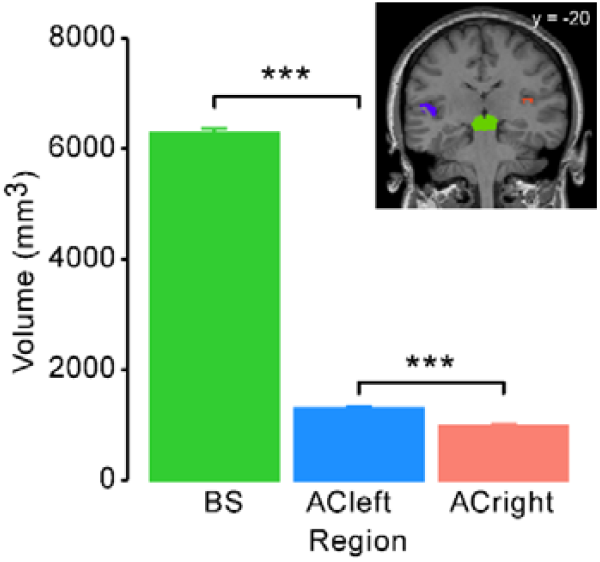
Volumetrics of the auditory brainstem-cortical pathway. BS, midbrain volume according to FreeSurfer brainstem parcellation [62]. AC, primary auditory cortex (Heschl’s gyrus) [60]. Auditory brainstem is larger than bilateral auditory cortex. AC size is larger in the left vs. right hemisphere. errorbars = ± 1 s.e.m., \*\*\**p* < 0.0001.

**Figure 5:**
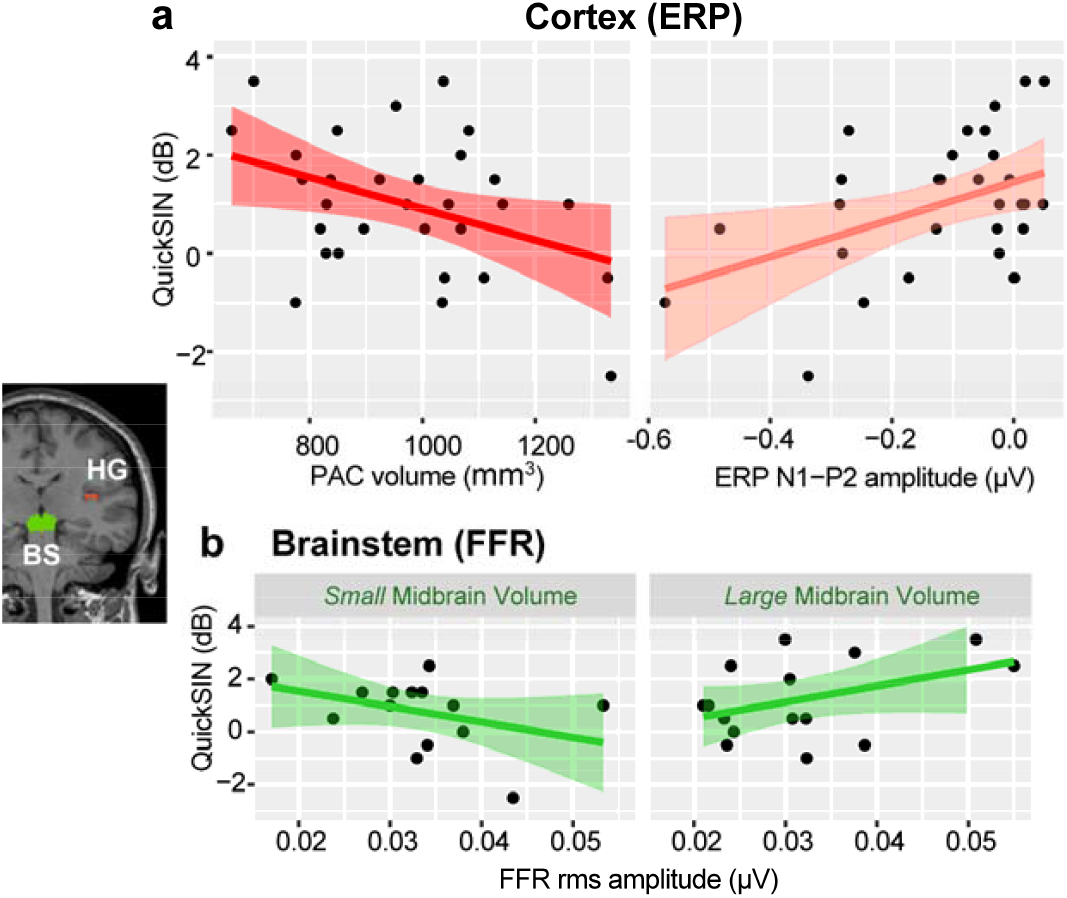
Structure-function-behavior relations underlying SIN processing. **(a)** Cortical data. Relations between behavioral QuickSIN performance and (i) PAC cortical volume and (ii) ERP amplitudes. Larger cortical anatomy and electrophysiological responses related to better SIN processing. (**b**) Brainstem data showing FFR-MRI interaction. For listeners with smaller midbrain volumes (lower 50^th^ percentile), stronger FFRs were associated with better QuickSIN scores. In contrast, in listeners with larger midbrains (upper 50^th^ percentile), stronger FFRs were associated with poorer QuickSIN. HG, Heschl’s gyrus (transverse temporal gyrus; primary auditory cortex). BS, brainstem (midbrain). shading = 95% CI.

## 4. Discussion

In this multimodal imaging study, we examined structure-function-behavior links between brainstem (FFR) and cortical (ERP) responses to noise-degraded speech and anatomical properties of their corresponding auditory midbrain-cortical structures. Our data reveal three key findings: (1) larger/thicker right (but not left) Heschl’s gyrus predicts better SIN listening performance; (2) HG morphology predicts the magnitude of listeners’ cortical response to speech (ERPs); and (3) a structure-function interaction at subcortical levels; listeners with smaller brainstem anatomy but larger speech FFRs show improved SIN perception.

Our cortical MRI volumetric data are consistent with previous reports showing larger density and leftward asymmetry of auditory cortex (HG) [80-82]. Schneider, *et al*. [40] reported HG volume ranging between ∼900-2000 mm^3^ depending on listeners’ musical training. While professional musicians have reliably larger (right) HG volumes than nonmusicians, amateur musicians (akin to the mean musicianship levels in our sample) do not differ from nonmusicians in aggregate HG volumetrics. Peelle, *et al*. [42] showed that in older adults, gray matter volume in *right* primary auditory cortex correlated with hearing acuity as measured by pure-tone audiometric thresholds; subtle hearing loss was associated with smaller gray matter volume. Importantly, control analyses confirmed hearing acuity was not associated with atrophy in motor regions, suggesting structure-function relationships were specific to auditory processing. We observe similar results for suprathreshold SIN perception.

Anatomical variability in auditory cortex has been related to several hearing behaviors including learning foreign speech sounds [84], tracking linguistic pitch patterns [85], melody discrimination [86], and speech [87] and musical sound [88] identification. In older adults, reduced temporal lobe volume correlates with poorer SIN performance as measured by the QuickSIN [89]. Our results broadly converge with these previous findings and demonstrate new links between auditory cortical morphology and complex hearing skills. We found that right transverse temporal gyrus morphology was related to improved (i.e., lower) QuickSIN scores (structure-behavior relation). In turn, larger ERP N1-P2 amplitudes to noise-degraded speech were associated with better QuickSIN (functional-behavior relation). That these brain-behavior correlations were right lateralized implies that more voluminous auditory cortical anatomy in the “non-linguistic” hemisphere supports better figure-ground speech perception. The critical role of right auditory-linguistic brain areas in SIN has been appreciated in recent functional EEG studies [90-93] and may reflect compensatory responses of the non-dominant hemisphere to help decipher degraded/ambiguous speech signals.

Correlations between speech FFRs and midbrain volumetrics revealed a critical interaction between the strength of electrophysiological responses and anatomical size. We found that in listeners with smaller midbrain volumes (bottom 50%-tile), stronger FFR responses were associated with *better* QuickSIN scores. On the contrary, in listeners with larger midbrains (upper 50%-tile), stronger FFRs were related with *worse* QuickSIN scores. These findings suggest that when the auditory brainstem is of lesser size, functional encoding of degraded speech is bolstered to provide additional compensatory processing. In contrast, when midbrain anatomy is already substantially large, further functional resources may not provide additional support to SIN perception—and may even reflect an inefficiency or redundancy in speech coding. Differences in neural organization, myelination, or synaptic density imply that larger morphology within the auditory pathways may not necessarily yield better computational efficiency as often assumed [i.e., “bigger is not better”; 94]. Alternatively, the FFR-MRI interactions observed here could highlight different listening strategies for SIN processing due to stronger anatomical support vs. functional encoding dependent on a certain listener’s profile [e.g., 88,95,96]. The data may also reflect quasi-ceiling/floor effects, if listeners with larger anatomy are already near ceiling in neural encoding and thus enjoy minimal added benefit from additional neural resources.

Our EEG-MRI findings have ramifications for understanding neuroplasticity observed in the auditory evoked potentials including the FFR. The strength to which brainstem responses capture voice pitch and harmonic timbre cues of complex signals relates to listeners’ perception of speech material [97; present study]. FFRs are also enhanced by various experiential factors including native language experience [98,99], music abilities [100,101], and perceptual learning [44,102]. FFR waveform enhancements are often interpreted to reflect plasticity in central auditory processing [29,31,99-101,103-107]. For example, several studies report that musicians have superior SIN processing and correspondingly more robust FFRs [29,33,108-110]. Yet, other studies have failed to find musician SIN advantages either at the behavioral [111-115] or FFR level [116]. Our data here demonstrate that the size of a listener’s midbrain anatomy interacts with the size of their functional FFR response and its correspondence with perceptual SIN abilities. Failures to replicate FFR-SIN effects may be related, at least in part, to unmeasured individual differences in midbrain anatomy [cf. 55]. This notion is supported by several MRI-M/EEG cortical findings showing larger HG morphology emits more robust auditory ERPs to speech and musical sounds [40,88]. We extend this prior work by demonstrating similar structure-function relations in subcortex between auditory midbrain anatomy and FFRs.

Physical factors of a listener are known to modulate auditory EEG responses. Head size (e.g., circumference) has a well-known influence on auditory brainstem response amplitude and latency [117-119]. Stronger FFRs to linguistic pitch stimuli have been observed in speakers of tonal languages vs. their English-speaking peers [98,106]. However, individuals of East Asian descent have different head sizes and shapes [120] compared to Caucasian listeners [cf. 121]. Insomuch as head size is related to the brain volumetrics assessed here, it is conceivable that differences in physical dimensions of the head could partly explain subtle group differences in FFRs reported previously. However, this account seems untenable by itself given that experience-dependent enhancements in the FFR are still observable among listeners of similar cultural background [100,122]. FFR neuroplasticity is also maintained even when controlling for confounding listener demographics and recording variables that can artificially inflate FFR amplitudes [55,101]. Accounts of our data due to simple size differences are also unlikely given the *interaction* we find between midbrain volumetrics, FFR magnitudes, and QuickSIN (Fig. 5b). Explanations based on mere EEG volume conduction principles [37,38] would, on the contrary, predict larger responses from larger anatomy across the board, which is not what we observe in our EEG-MRI data.

Instead, our brainstem findings are most parsimoniously explained as a layering of functional processing (as index via FFR) that can override fixed, structural predispositions in midbrain anatomy depending on the perceptual demands of speech processing [see also 123]. Of note, our FFRs/ERPs were also recorded during active SIN perception tasks. It is possible the observed structure-function-behavior interactions of the auditory brainstem might only emerge under states of active (attentional) and not passive listening conditions [e.g., 18]. Future studies are needed to test this possibility along with whether anatomical variation in brainstem anatomy moderates or mediates experience-dependent [98,100,101,106] and learning-related plasticity in the FFR [44,102,124].

## Author Contributions

Conceptualization, G.B.; data collection, R.R., J.S., J. M., formal analysis, G.B.; validation, H.C., writing, all authors; funding acquisition, G.B. All authors have read and agreed to the published version of the manuscript.

## Institutional Review Board Statement

The study was conducted in accordance with the Declaration of Helsinki and protocols approved by the Indiana University Institutional Review Board (MRI: #15650, approved 7/15/22; EEG: #14860, approved 4/7/22).

## Informed Consent Statement

Written informed consent was obtained from all participants involved in the study.

## Data Availability Statement

The data presented in this study are not publicly available due to privacy or ethical restrictions but are available on request from the corresponding author.

## Footnotes

1 EEG recording originally included 64 channels positioned at 10-10 scalp locations. Blink artifacts were correctedin the continuous EEGs topographically using principal component analysis (PCA) [52]. For the purposes of this study, we analyzed FFRs/ERPs isolated to the Cz electrode were FFRs and ERPs are maximal on the scalp [26,53]. Full multichannel results of the electrophysiological data and related behavioral task will be reported elsewhere.

2 Note that thickness measures are not relevant to the FreeSurfer brainstem segmentation pipeline [62]. Thus, only volume could be directly compared across anatomical levels.

3 We only considered right auditory cortex in this analysis since left hemisphere was not related to QuickSIN (see Fig. 3). Similarly, only FFRs/ERPs to noise-degraded speech were considered as we were interested in assessing brain-behavior relations underlying SIN processing rather than clean speech.

## References

1. Bidelman, G.M. Communicating in challenging environments: Noise and reverberation. In Springer Handbook of Auditory Research: The frequency-following response: A window into human communication, Kraus, N., Anderson, S., White-Schwoch, T., Fay, R.R., Popper, A.N., Eds.; Springer Nature: New York, N.Y., 2017.

2. Anderson, S.; Parbery-Clark, A.; Yi, H.G.; Kraus, N. A neural basis of speech-in-noise perception in older adults. Ear Hear. 2011, 32, 750–757. doi:10.1097/AUD.0b013e31822229d3.

3. Song, J.H.; Skoe, E.; Banai, K.; Kraus, N. Perception of speech in noise: Neural correlates. J. Cogn. Neurosci. 2011, 23, 2268–2279. doi:10.1162/jocn.2010.21556.

4. Bidelman, G.M.; Momtaz, S. Subcortical rather than cortical sources of the frequency-following response (FFR) relate to speech-in-noise perception in normal-hearing listeners. Neurosci. Lett. 2021, 746, 135664. doi:10.1016/j.neulet.2021.135664.

5. Bidelman, G.M.; Davis, M.K.; Pridgen, M.H. Brainstem-cortical functional connectivity for speech is differentially challenged by noise and reverberation. Hear. Res. 2018, 367, 149–160. doi:10.1016/j.heares.2018.05.018.

6. Billings, C.J.; McMillan, G.P.; Penman, T.M.; Gille, S.M. Predicting perception in noise using cortical auditory evoked potentials. Journal of the Association for Research in Otolaryngology 2013, 14, 891– 903. doi:10.1007/s10162-013-0415-y.

7. Garrett, M.F.; Vasilkov, V.; Mauermann, M.; Devolder, P.; Wilson, J.L.; Gonzales, L.; Henry, K.S.; Verhulst, S. Deciphering compromised speech-in-noise intelligibility in older listeners: The role of cochlear synaptopathy. eneuro 2025, 12, ENEURO.0182–0124.2024. doi:10.1523/eneuro.0182-24.2024.

8. Lagacé, J.; Jutras, B.; Gagné, J.P. Auditory processing disorder and speech perception problems in noise: finding the underlying origin. Am J Audiol 2010, 19, 17–25. doi:10.1044/1059-0889(2010/09-0022).

9. Cunningham, J.; Nicol, T.; Zecker, S.G.; Bradlow, A.; Kraus, N. Neurobiologic responses to speech in noise in children with learning problems: Deficits and strategies for improvement. Clin. Neurophysiol. 2001, 112, 758–767. doi:S1388-2457(01)00465-5 [pii].

10. Warrier, C.M.; Johnson, K.L.; Hayes, E.A.; Nicol, T.; Kraus, N. Learning impaired children exhibit timing deficits and training-related improvements in auditory cortical responses to speech in noise. Exp. Brain Res. 2004, 157, 431–441. doi:10.1007/s00221-004-1857-6.

11. Putter-Katz, H.; Adi-Bensaid, L.; Feldman, I.; Hildesheimer, M. Effects of speech in noise and dichotic listening intervention programs on central auditory processing disorders. J. Basic Clin. Physiol. Pharmacol. 2008, 19, 301–316. doi:10.1515/jbcpp.2008.19.3-4.301.

12. Dole, M.; Meunier, F.; Hoen, M. Functional correlates o fthes peech-in-noise perception impairment in dyslexia: An MRI study. Neuropsychologia 2014, 60, 103–114.

13. Dole, M.; Hoen, M.; Meunier, F. Speech-in-noise perception deficit in adults with dyslexia: effects of background type and listening configuration. Neuropsychologia 2012, 50, 1543–1552. doi:10.1016/j.neuropsychologia.2012.03.007.

14. Middelweerd, M.J.; Festen, J.M.; Plomp, R. Difficulties with speech intelligibility in noise in spite of a normal pure-tone audiogram. Audiology 1990, 29, 1–7.

15. Bidelman, G.M.; Howell, M. Functional changes in inter- and intra-hemispheric cortical processing underlying degraded speech perception. Neuroimage 2016, 124, 581–590. doi:10.1016/j.neuroimage.2015.09.020.

16. Fitzgerald, M.B.; Ward, K.M.; Gianakas, S.P.; Smith, M.L.; Blevins, N.H.; Swanson, A.P. Speech-in-Noise Assessment in the Routine Audiologic Test Battery: Relationship to Perceived Auditory Disability. Ear Hear. 2024, 45, 816–826. doi:10.1097/aud.0000000000001472.

17. Fitzgerald, M.B.; Gianakas, P.; Qian, Z.J.; Losorelli, S.; Swanson, A.C. Preliminary guidelines for replacing word-recognition in quiet with speech in noise assessment in the routine audiologic test battery. Ear Hear. 2023.

18. Price, C.N.; Bidelman, G.M. Attention reinforces human corticofugal system to aid speech perception in noise. Neuroimage 2021, 235, 118014. doi:10.1016/j.neuroimage.2021.118014.

19. Bidelman, G.M.; Moreno, S.; Alain, C. Tracing the emergence of categorical speech perception in the human auditory system. Neuroimage 2013, 79, 201–212. doi:10.1016/j.neuroimage.2013.04.093.

20. Picton, T.W.; Alain, C.; Woods, D.L.; John, M.S.; Scherg, M.; Valdes-Sosa, P.; Bosch-Bayard, J.; Trujillo, N.J. Intracerebral sources of human auditory-evoked potentials. Audiol. Neurootol. 1999, 4, 64– 79. doi:13823 [pii].

21. Krishnan, A. Human frequency following response. In Auditory evoked potentials: Basic principles and clinical application, Burkard, R.F., Don, M., Eggermont, J.J., Eds.; Lippincott Williams & Wilkins: Baltimore, 2007; pp. 313–335.

22. Skoe, E.; Kraus, N. Auditory brain stem response to complex sounds: A tutorial. Ear Hear. 2010, 31, 302–324. doi:10.1097/AUD.0b013e3181cdb272.

23. Bidelman, G.M. Subcortical sources dominate the neuroelectric auditory frequency-following response to speech. Neuroimage 2018, 175, 56–69.

24. Gorina-Careta, N.; Kurkela, J.L.O.; Hämäläinen, J.; Astikainen, P.; Escera, C. Neural generators of the frequency-following response elicited to stimuli of low and high frequency: A magnetoencephalographic (MEG) study. Neuroimage 2021, 231, 117866. doi:10.1016/j.neuroimage.2021.117866.

25. Coffey, E.B.J.; Nicol, T.; White-Schwoch, T.; Chandrasekaran, B.; Krizman, J.; Skoe, E.; Zatorre, R.J.; Kraus, N. Evolving perspectives on the sources of the frequency-following response. Nature Communications 2019, 10, 5036. doi:10.1038/s41467-019-13003-w.

26. Bidelman, G.M. Multichannel recordings of the human brainstem frequency-following response: Scalp topography, source generators, and distinctions from the transient ABR. Hear. Res. 2015, 323, 68–80. doi:S0378-5955(15)00022-2 [pii] 10.1016/j.heares.2015.01.011.

27. Ross, B.; Tremblay, K.L.; Alain, C. Simultaneous EEG and MEG recordings reveal vocal pitch elicited cortical gamma oscillations in young and older adults. Neuroimage 2020, 204, 116253. doi:10.1016/j.neuroimage.2019.116253.

28. López-Caballero, F.; Martin-Trias, P.; Ribas-Prats, T.; Gorina-Careta, N.; Bartrés-Faz, D.; Escera, C. Effects of cTBS on the frequency-following response and other auditory evoked potentials. Front. Hum. Neurosci. 2020, 14. doi:10.3389/fnhum.2020.00250.

29. Parbery-Clark, A.; Skoe, E.; Kraus, N. Musical experience limits the degradative effects of background noise on the neural processing of sound. J. Neurosci. 2009, 29, 14100–14107. doi:29/45/14100 [pii] 10.1523/JNEUROSCI.3256-09.2009.

30. Anderson, S.; Skoe, E.; Chandrasekaran, B.; Kraus, N. Neural timing is linked to speech perception in noise. J. Neurosci. 2010, 30, 4922–4926.

31. Coffey, E.B.J.; Chepesiuk, A.M.P.; Herholz, S.C.; Baillet, S.; Zatorre, R.J. Neural correlates of early sound encoding and their relationship to speech-in-noise perception. Front. Neurosci. 2017, 11. doi:10.3389/fnins.2017.00479.

32. Parbery-Clark, A.; Marmel, F.; Bair, J.; Kraus, N. What subcortical-cortical relationships tell us about processing speech in noise. Eur. J. Neurosci. 2011, 33, 549–557. doi:10.1111/j.1460-9568.2010.07546.x.

33. Bidelman, G.M.; Krishnan, A. Effects of reverberation on brainstem representation of speech in musicians and non-musicians. Brain Res. 2010, 1355, 112–125.

34. Yellamsetty, A.; Bidelman, G.M. Brainstem correlates of concurrent speech identification in adverse listening conditions. Brain Res. 2019, 1714, 182–192. doi:10.1016/j.brainres.2019.02.025.

35. Saiz-Alía, M.; Forte, A.E.; Reichenbach, T. Individual differences in the attentional modulation of the human auditory brainstem response to speech inform on speech-in-noise deficits. Sci. Rep. 2019, 9, 14131. doi:10.1038/s41598-019-50773-1.

36. Skoe, E.; Krizman, J.; Spitzer, E.; Kraus, N. The auditory brainstem is a barometer of rapid auditory learning. Neuroscience 2013, 243, 104–114. doi:S0306-4522(13)00224-8 [pii] 10.1016/j.neuroscience.2013.03.009.

37. Scherg, M. Fundamentals of dipole source potential analysis. In: Auditory evoked magnetic fields and electric potentials. In Advances in Audiology, Grandori, F., Hoke, M., Romani, G.L., Eds.; Karger: Basel, 1990; pp. 40–69.

38. Scherg, M.; Berg, P.; Nakasato, N.; Beniczky, S. Taking the EEG back into the brain: The power of multiple discrete sources. Front. Neurol. 2019, 10. doi:10.3389/fneur.2019.00855.

39. Schneider, P.; Andermann, M.; Wengenroth, M.; Goebel, R.; Flor, H.; Rupp, A.; Diesch, E. Reduced volume of Heschl’s gyrus in tinnitus. Neuroimage 2009, 45, 927–939. doi:10.1016/j.neuroimage.2008.12.045.

40. Schneider, P.; Scherg, M.; Dosch, H.G.; Specht, H.J.; Gutschalk, A.; Rupp, A. Morphology of Heschl’s gyrus reflects enhanced activation in the auditory cortex of musicians. Nat. Neurosci. 2002, 5, 688–694. doi:10.1038/nn871nn871 [pii].

41. Wengenroth, M.; Blatow, M.; Heinecke, A.; Reinhardt, J.; Stippich, C.; Hofmann, E.; Schneider, P. Increased volume and function of right auditory cortex as a marker for absolute pitch. Cereb. Cortex 2013, 24, 1127–1137. doi:10.1093/cercor/bhs391.

42. Peelle, J.E.; Troiani, V.; Grossman, M.; Wingfield, A. Hearing loss in older adults affects neural systems supporting speech comprehension. J. Neurosci. 2011, 31, 12638–12643.

43. Coffey, E.B.J.; Musacchia, G.; Zatorre, R.J. Cortical correlates of the auditory frequency-following and onset responses: EEG and fMRI evidence. J. Neurosci. 2017, 37, 830–838. doi:10.1523/jneurosci.1265-16.2016.

44. Chandrasekaran, B.; Kraus, N.; Wong, P.C. Human inferior colliculus activity relates to individual differences in spoken language learning. J. Neurophysiol. 2012, 107, 1325–1336. doi:jn.00923.2011 [pii] 10.1152/jn.00923.2011.

45. Oldfield, R.C. The assessment and analysis of handedness: The Edinburgh inventory. Neuropsychologia 1971, 9, 97–113.

46. Bidelman, G.M. Subcortical sources dominate the neuroelectric auditory frequency-following response to speech. Neuroimage 2018, 175, 56–69. doi:10.1016/j.neuroimage.2018.03.060.

47. Coffey, E.B.; Herholz, S.C.; Chepesiuk, A.M.; Baillet, S.; Zatorre, R.J. Cortical contributions to the auditory frequency-following response revealed by MEG. Nat Commun 2016, 7, 11070. doi:10.1038/ncomms11070.

48. Bidelman, G.M.; Weiss, M.W.; Moreno, S.; Alain, C. Coordinated plasticity in brainstem and auditory cortex contributes to enhanced categorical speech perception in musicians. Eur. J. Neurosci. 2014, 40, 2662–2673.

49. Bidelman, G.M.; Alain, C. Musical training orchestrates coordinated neuroplasticity in auditory brainstem and cortex to counteract age-related declines in categorical vowel perception. J. Neurosci. 2015, 35, 1240 –1249.

50. Bidelman, G.M. Towards an optimal paradigm for simultaneously recording cortical and brainstem auditory evoked potentials. J Neurosci Methods 2015, 241, 94–100. doi:10.1016/j.jneumeth.2014.12.019.

51. Musacchia, G.; Strait, D.; Kraus, N. Relationships between behavior, brainstem and cortical encoding of seen and heard speech in musicians and non-musicians. Hear. Res. 2008, 241, 34–42.

52. Picton, T.W.; van Roon, P.; Armilio, M.L.; Berg, P.; Ille, N.; Scherg, M. The correction of ocular artifacts: A topographic perspective. Clin. Neurophysiol. 2000, 111, 53–65.

53. Bidelman, G.M.; Bush, L.C.; Boudreaux, A.M. Effects of noise on the behavioral and neural categorization of speech. Front. Neurosci. 2020, 14, 1–13. doi:10.3389/fnins.2020.00153.

54. Bidelman, G.M.; Mahmud, M.S.; Yeasin, M.; Shen, D.; Arnott, S.; Alain, C. Age-related hearing loss increases full-brain connectivity while reversing directed signaling within the dorsal-ventral pathway for speech. Brain Structure and Function 2019, 224, 2661–2676.

55. Bidelman, G.M.; Sisson, A.; Rizzi, R.; MacLean, J.; Baer, K. Myogenic artifacts masquerade as neuroplasticity in the auditory frequency-following response. Front. Neurosci. 2024, 18, 1–13. doi:10.3389/fnins.2024.1422903.

56. Levitas, D.; Hayashi, S.; Vinci-Booher, S.; Heinsfeld, A.; Bhatia, D.; Lee, N.; Galassi, A.; Niso, G.; Pestilli, F. ezBIDS: Guided standardization of neuroimaging data interoperable with major data archives and platforms. Scientific Data 2024, 11, 179. doi:10.1038/s41597-024-02959-0.

57. Fischl, B. FreeSurfer. Neuroimage 2012, 62, 774–781. doi:10.1016/j.neuroimage.2012.01.021.

58. Reuter, M.; Rosas, H.D.; Fischl, B. Highly accurate inverse consistent registration: a robust approach. Neuroimage 2010, 53, 1181–1196. doi:10.1016/j.neuroimage.2010.07.020.

59. Fischl, B.; van der Kouwe, A.; Destrieux, C.; Halgren, E.; Segonne, F.; Salat, D.H.; Busa, E.; Seidman, L.J.; Goldstein, J.; Kennedy, D.; et al. Automatically parcellating the human cerebral cortex. Cereberal Cortex 2004, 14, 11–22.

60. Desikan, R.S.; Segonne, F.; Fischl, B.; Quinn, B.T.; Dickerson, B.C.; Blacker, D.; Buckner, R.L.; Dale, A.M.; Maguire, R.P.; Hyman, B.T.; et al. An automated labeling system for subdividing the human cerebral cortex on MRI scans into gyral based regions of interest. Neuroimage 2006, 31, 968–980. doi:10.1016/j.neuroimage.2006.01.021.

61. Dale, A.M.; Fischl, B.; Sereno, M.I. Cortical surface-based analysis. I. Segmentation and surface reconstruction. Neuroimage 1999, 9, 179–194. doi:10.1006/nimg.1998.0395.

62. Iglesias, J.E.; Van Leemput, K.; Bhatt, P.; Casillas, C.; Dutt, S.; Schuff, N.; Truran-Sacrey, D.; Boxer, A.; Fischl, B. Bayesian segmentation of brainstem structures in MRI. Neuroimage 2015, 113, 184–195. doi:10.1016/j.neuroimage.2015.02.065.

63. Fischl, B.; Dale, A.M. Measuring the thickness of the human cerebral cortex from magnetic resonance images. Proc. Natl. Acad. Sci. U. S. A. 2000, 97, 11050–11055. doi:10.1073/pnas.200033797.

64. Fischl, B.; Sereno, M.I.; Dale, A.M. Cortical surface-based analysis. II: Inflation, flattening, and a surface-based coordinate system. Neuroimage 1999, 9, 195–207. doi:10.1006/nimg.1998.0396.

65. Reuter, M.; Schmansky, N.J.; Rosas, H.D.; Fischl, B. Within-subject template estimation for unbiased longitudinal image analysis. Neuroimage 2012, 61, 1402–1418. doi:10.1016/j.neuroimage.2012.02.084.

66. Han, X.; Jovicich, J.; Salat, D.; van der Kouwe, A.; Quinn, B.; Czanner, S.; Busa, E.; Pacheco, J.; Albert, M.; Killiany, R.; et al. Reliability of MRI-derived measurements of human cerebral cortical thickness: the effects of field strength, scanner upgrade and manufacturer. Neuroimage 2006, 32, 180–194. doi:10.1016/j.neuroimage.2006.02.051.

67. Fischl, B.; Sereno, M.I.; Tootell, R.B.H.; Dale, A.M. High-resolution intersubject averaging and a coordinate system for the cortical surface. Hum. Brain Mapp. 1999, 8, 272–284. doi: https://doi.org/10.1002/(SICI)1097-0193(1999)8:4<272::AID-HBM10>3.0.CO;2-4.

68. Killion, M.C.; Niquette, P.A.; Gudmundsen, G.I.; Revit, L.J.; Banerjee, S. Development of a quick speech-in-noise test for measuring signal-to-noise ratio loss in normal-hearing and hearing-impaired listeners. J. Acoust. Soc. Am. 2004, 116, 2395–2405.

69. Bidelman, G.M.; Yoo, J. Musicians show improved speech segregation in competitive, Multi-talker cocktail party scenarios. Front Psychol 2020, 11, 1927. doi:10.3389/fpsyg.2020.01927.

70. Bidelman, G.M.; Mahmud, M.S.; Yeasin, M.; Shen, D.; Arnott, S.R.; Alain, C. Age-related hearing loss increases full-brain connectivity while reversing directed signaling within the dorsal-ventral pathway for speech. Brain Struct Funct 2019, 224, 2661–2676. doi:10.1007/s00429-019-01922-9.

71. R-Core-Team R: A language and environment for statistical computing, R Foundation for Statistical Computing, Vienna, Austria. URL https://www.R-project.org/: 2020.

72. Schaefer, T.; Ecker, C. fsbrain: An R package for the visualization of structural neuroimaging data. bioRxiv [pre-print]. doi:10.1101/2020.09.18.302935 2020.

73. Alain, C.; Snyder, J.S.; He, Y.; Reinke, K.S. Changes in auditory cortex parallel rapid perceptual learning. Cereb. Cortex 2007, 17, 1074–1084. doi:bhl018 [pii] 10.1093/cercor/bhl018.

74. MacLean, J.; Stirn, J.; Sisson, A.; Bidelman, G.M. Short- and long-term neuroplasticity interact during the perceptual learning of concurrent speech. Cereb. Cortex 2024, 34, 1–13. doi:10.1093/cercor/bhad543.

75. Bidelman, G.; Powers, L. Response properties of the human frequency-following response (FFR) to speech and non-speech sounds: level dependence, adaptation and phase-locking limits. Int. J. Audiol. 2018, 57, 665–672. doi:10.1080/14992027.2018.1470338.

76. Bidelman, G.M.; Price, C.N.; Shen, D.; Arnott, S.R.; Alain, C. Afferent-efferent connectivity between auditory brainstem and cortex accounts for poorer speech-in-noise comprehension in older adults. Hear. Res. 2019, 382, 1–12. doi:10.1016/j.heares.2019.107795.

77. Presacco, A.; Simon, J.Z.; Anderson, S. Evidence of degraded representation of speech in noise, in the aging midbrain and cortex. J. Neurophysiol. 2016, 116, 2346–2355. doi:10.1152/jn.00372.2016.

78. Billings, C.J.; Bennett, K.O.; Molis, M.R.; Leek, M.R. Cortical encoding of signals in noise: Effects of stimulus type and recording paradigm. Ear Hear. 2010, 32, 53–60.

79. Lai, J.; Alain, C.; Bidelman, G.M. Cortical-brainstem interplay during speech perception in older adults with and without hearing loss. Front. Neurosci. 2023, 17, 1–12. doi:10.3389/fnins.2023.1075368.

80. Schneider, P.; Sluming, V.; Roberts, N.; Scherg, M.; Goebel, R.; Specht, H.J.; Dosch, H.G.; Bleeck, S.; Stippich, C.; Rupp, A. Structural and functional asymmetry of lateral Heschl’s gyrus reflects pitch perception preference. Nat. Neurosci. 2005, 8, 1241–1247. doi:10.1038/nn1530.

81. Jancke, L.; Steinmetz, H. Anatomical brain asymmetries and their relevance for functional asymmetries. In The Asymmetrical Brain, Davidson, R.J., Hugdahl, K., Eds.; MIT Press: Boston, MA, 2003; pp. 187– 229.

82. Hutsler, J.J. The specialized structure of human language cortex: pyramidal cell size asymmetries within auditory and language-associated regions of the temporal lobes. Brain Lang. 2003, 86, 226–242. doi:10.1016/s0093-934x(02)00531-x.

83. Du, Y.; Buchsbaum, B.R.; Grady, C.L.; Alain, C. Noise differentially impacts phoneme representations in the auditory and speech motor systems. Proc. Natl. Acad. Sci. U. S. A. 2014, 111, 1–6.

84. Golestani, N.; Molko, N.; Dehaene, S.; LeBihan, D.; Pallier, C. Brain structure predicts the learning of foreign speech sounds. Cerebral Cortex 2007, 17, 575–582. doi:10.1093/cercor/bhk001.

85. Wong, P.C.; Warrier, C.M.; Penhune, V.B.; Roy, A.K.; Sadehh, A.; Parrish, T.B.; Zatorre, R.J. Volume of left Heschl’s Gyrus and linguistic pitch learning. Cereb. Cortex 2008, 18, 828–836. doi:bhm115 [pii] 10.1093/cercor/bhm115.

86. Foster, N.E.; Zatorre, R.J. Cortical structure predicts success in performing musical transformation judgments. Neuroimage 2010, 53, 26–36. doi:S1053-8119(10)00886-4 [pii] 10.1016/j.neuroimage.2010.06.042.

87. Fuhrmeister, P.; Myers, E.B. Structural neural correlates of individual differences in categorical perception. Brain and Language 2021, 215. doi:10.1016/j.bandl.2021.104919.

88. Mankel, K.; Shrestha, U.; Tipirneni-Sajja, A.; Bidelman, G.M. Functional plasticity coupled with structural predispositions in auditory cortex shape successful music category learning. Front. Neurosci. 2022, 16, 1–14.

89. Jiang, K.; Albert, M.S.; Coresh, J.; Couper, D.J.; Gottesman, R.F.; Hayden, K.M.; Jack, C.R., Jr.; Knopman, D.S.; Mosley, T.H.; Pankow, J.S.; et al. Cross-Sectional Associations of Peripheral Hearing, Brain Imaging, and Cognitive Performance With Speech-in-Noise Performance: The Aging and Cognitive Health Evaluation in Elders Brain Magnetic Resonance Imaging Ancillary Study. Am J Audiol 2024, 33, 683–694. doi:10.1044/2024_aja-23-00108.

90. Carter, J.A.; Bidelman, G.M. Auditory cortex is susceptible to lexical influence as revealed by informational vs. energetic masking of speech categorization. Brain Res. 2021, 1759, 147385. doi:10.1101/2020.10.20.347724.

91. Bidelman, G.M.; Howell, M. Functional changes in inter- and intra-hemispheric auditory cortical processing underlying degraded speech perception. Neuroimage 2016, 124, 581–590.

92. Price, C.N.; Alain, C.; Bidelman, G.M. Auditory-frontal channeling in α and β bands is altered by age-related hearing loss and relates to speech perception in noise. Neuroscience 2019, 423, 18–28. doi:10.1016/j.neuroscience.2019.10.044.

93. Mahmud, S.; Ahmed, F.; Al-Fahad, R.; Moinuddin, K.A.; Yeasin, M.; Alain, C.; Bidelman, G.M. Decoding hearing-related changes in older adults’ spatiotemporal neural processing of speech using machine learning. Front. Neurosci. 2020, 14, 1–15. doi:10.1101/786566.

94. Kanai, R.; Rees, G. The structural basis of inter-individual differences in human behaviour and cognition. Nature Reviews Neuroscience 2011, 12, 231–242. doi:10.1038/nrn3000.

95. Rizzi, R.; Bidelman, G.M. Functional benefits of continuous vs. categorical listening strategies on the neural encoding and perception of noise-degraded speech. Brain Res. 2024, 1844, 1–12. doi:10.1016/j.brainres.2024.149166.

96. Bidelman, G.M.; Walker, B.S. Plasticity in auditory categorization is supported by differential engagement of the auditory-linguistic network. Neuroimage 2019, 201, 1–10. doi:10.1016/j.neuroimage.2019.116022.

97. Weiss, M.W.; Bidelman, G.M. Listening to the brainstem: Musicianship enhances intelligibility of subcortical representations for speech. J. Neurosci. 2015, 35, 1687–1691.

98. Krishnan, A.; Gandour, J.T.; Bidelman, G.M. The effects of tone language experience on pitch processing in the brainstem. J. Neurolinguistics 2010, 23, 81–95.

99. Zhao, T.C.; Kuhl, P.K. Linguistic effect on speech perception observed at the brainstem. Proc. Natl. Acad. Sci. U. S. A. 2018, 115, 8716–8721. doi:10.1073/pnas.1800186115.

100. Wong, P.C.; Skoe, E.; Russo, N.M.; Dees, T.; Kraus, N. Musical experience shapes human brainstem encoding of linguistic pitch patterns. Nat. Neurosci. 2007, 10, 420–422. doi:10.1038/nn1872.

101. Mankel, K.; Bidelman, G.M. Inherent auditory skills rather than formal music training shape the neural encoding of speech. Proc. Natl. Acad. Sci. U. S. A. 2018, 115, 13129–13134. doi:10.1073/pnas.1811793115.

102. Reetzke, R.; Xie, Z.; Llanos, F.; Chandrasekaran, B. Tracing the trajectory of sensory plasticity across different stages of speech learning in adulthood. Curr. Biol. 2018, 28, 1419–1427.e1414. doi:10.1016/j.cub.2018.03.026.

103. Kraus, N.; Skoe, E.; Parbery-Clark, A.; Ashley, R. Experience-induced malleability in neural encoding of pitch, timbre, and timing. Ann. N. Y. Acad. Sci. 2009, 1169, 543–557. doi:NYAS04549 [pii] 10.1111/j.1749-6632.2009.04549.x.

104. Musacchia, G.; Sams, M.; Skoe, E.; Kraus, N. Musicians have enhanced subcortical auditory and audiovisual processing of speech and music. Proc. Natl. Acad. Sci. U. S. A. 2007, 104, 15894–15898. doi:10.1073/pnas.0701498104.

105. Krizman, J.; Marian, V.; Shook, A.; Skoe, E.; Kraus, N. Subcortical encoding of sound is enhanced in bilinguals and relates to executive function advantages. Proc. Natl. Acad. Sci. U. S. A. 2012, 109, 7877– 7881. doi:10.1073/pnas.1201575109.

106. Krishnan, A.; Xu, Y.; Gandour, J.T.; Cariani, P. Encoding of pitch in the human brainstem is sensitive to language experience. Brain Research Cognitive Brain Research 2005, 25, 161–168.

107. Kraus, N.; Slater, J.; Thompson, E.C.; Hornickel, J.; Strait, D.L.; Nicol, T.; White-Schwoch, T. Music enrichment programs improve the neural encoding of speech in at-risk children. J. Neurosci. 2014, 34, 11913–11918. doi:10.1523/jneurosci.1881-14.2014.

108. Hennessy, S.; Mack, W.J.; Habibi, A. Speech-in-noise perception in musicians and non-musicians: A multi-level meta-analysis. Hear. Res. 2022, 108442. doi:10.1016/j.heares.2022.108442.

109. Bidelman, G.M.; Brown, J.A.; Rizzi, R.; MacLean, J. Neuroplastic effects of music expertise on speech-language processing. In The Cambridge Handbook of Language and Brain, Andrews, E., Kiran, S., Eds.; Cambridge University Press: Cambridge, UK, 2025; pp. 423–458.

110. Coffey, E.B.J.; Mogilever, N.B.; Zatorre, R.J. Speech-in-noise perception in musicians: A review. Hear. Res. 2017, 352, 49–69. doi:10.1016/j.heares.2017.02.006.

111. Boebinger, D.; Evans, S.; Rosen, S.; Lima, C.F.; Manly, T.; Scott, S.K. Musicians and non-musicians are equally adept at perceiving masked speech. J. Acoust. Soc. Am. 2015, 137, 378–387. doi:10.1121/1.4904537.

112. Madsen, S.M.K.; Whiteford, K.L.; Oxenham, A.J. Musicians do not benefit from differences in fundamental frequency when listening to speech in competing speech backgrounds. Sci. Rep. 2017, 7, 12624. doi:10.1038/s41598-017-12937-9.

113. Ruggles, D.R.; Freyman, R.L.; Oxenham, A.J. Influence of musical training on understanding voiced and whispered speech in noise. PLoS One 2014, 9, e86980.

114. Yeend, I.; Beach, E.F.; Sharma, M.; Dillon, H. The effects of noise exposure and musical training on suprathreshold auditory processing and speech perception in noise. Hear. Res. 2017, 353, 224–236. doi:10.1016/j.heares.2017.07.006.

115. Escobar, J.; Mussoi, B.S.; Silberer, A.B. The effect of musical training and working memory in adverse listening situations. Ear Hear. 2020, 41, 278–288. doi:10.1097/aud.0000000000000754.

116. Whiteford, K.L.; Baltzell, L.S.; Chiu, M.; Cooper, J.K.; Faucher, S.; Goh, P.Y.; Hagedorn, A.; Irsik, V.C.; Irvine, A.; Lim, S.-J.; et al. Large-scale multi-site study shows no association between musical training and early auditory neural sound encoding. Nature Communications 2025, 16, 7152. doi:10.1038/s41467-025-62155-5.

117. Stockard, J.J.; Stockard, J.E.; Sharbrough, F.W. Nonpathologic factors influencing brainstem auditory evoked potentials. Am. J. EEG Technol. 1978, 18, 177–209. doi:10.1080/00029238.1978.11106793.

118. Trune, D.R.; Mitchell, C.; Phillips, D.S. The relative importance of head size, gender and age on the auditory brainstem response. Hear. Res. 1988, 32, 165–174. doi:10.1016/0378-5955(88)90088-3.

119. Chambers, R.D.; Matthies, M.L.; Griffiths, S.K. Correlations between various measures of head size and auditory brainstem response latencies. Hear. Res. 1989, 41, 179–187. doi:10.1016/0378-5955(89)90009-9.

120. Ball, R.; Shu, C.; Xi, P.; Rioux, M.; Luximon, Y.; Molenbroek, J. A comparison between Chinese and Caucasian head shapes. Appl. Ergon. 2010, 41, 832–839. doi:10.1016/j.apergo.2010.02.002.

121. Ulehlová, L.; Voldrich, L.; Janisch, R. Correlative study of sensory cell density and cochlear length in humans. Hear. Res. 1987, 28, 149–151. doi:10.1016/0378-5955(87)90045-1.

122. Bidelman, G.M.; Gandour, J.T.; Krishnan, A. Cross-domain effects of music and language experience on the representation of pitch in the human auditory brainstem. J. Cogn. Neurosci. 2011, 23, 425–434. doi:10.1162/jocn.2009.21362.

123. Krishnan, A.; Gandour, J.T.; Ananthakrishnan, S.; Bidelman, G.M.; Smalt, C.J. Functional ear (a)symmetry in brainstem neural activity relevant to encoding of voice pitch: A precursor for hemispheric specialization? Brain Lang. 2011, 119, 226–231. doi:S0093-934X(11)00100-3 [pii] 10.1016/j.bandl.2011.05.001.

124. Carcagno, S.; Plack, C.J. Subcortical plasticity following perceptual learning in a pitch discrimination task. Journal of the Association for Research in Otolaryngology 2011, 12, 89–100. doi:10.1007/s10162-010-0236-1.

